# Comparative analysis of genetic diversity and differentiation of cauliflower (*Brassica oleracea* var. *botrytis*) accessions from two *ex situ* genebanks

**DOI:** 10.1101/238840

**Authors:** Eltohamy A. A. Yousef, Thomas Müller, Andreas Börner, Karl J. Schmid

**Affiliations:** Department of Crop Biodiversity and Breeding Informatics, Faculty of Agriculture, University of Hohenheim, Stuttgart, Germany; Department of Horticulture, Faculty of Agriculture, University of Suez Canal, Ismailia, Egypt; Research (IPK), Seeland OT Gatersleben, Germany

## Abstract

Cauliflower (*Brassica oleracea* var. *botrytis*) is an important vegetable crop for human nutrition. We characterized 192 cauliflower accessions from the USDA and IPK genebanks with genotyping by sequencing (GBS). They originated from 26 different countries and represent about 44% of all cauliflower accessions in both genebanks. The analysis of genetic diversity revealed that accessions formed two major groups that represented the two genebanks and were not related to the country of origin. This differentiation was robust with respect to the analysis methods that included principal component analysis, ADMIXTURE and neighbor-joining trees. Genetic diversity was higher in the USDA collection and significant phenotypic differences between the two genebanks were found in three out of six traits investigated. GBS data have a high proportion of missing data, but we observed that the exclusion of single nucleotide polymorphisms (SNPs) with missing data or the imputation of missing SNP alleles produced very similar results. The results indicate that the composition and type of accessions have a strong effect on the structure of genetic diversity of *ex situ* collections, although regeneration procedures and local adaptation to regeneration conditions may also contribute to a divergence. *F_st_*-based outlier tests of genetic differentiation identified only a small proportion (<1%) of SNPs that are highly differentiated between the two genebanks, which indicates that selection during seed regeneration is not a major cause of differentiation between genebanks. Seed regeneration procedures of both genebanks do not result in different levels of genetic drift and loss of genetic variation. We therefore conclude that the composition and type of accessions mainly influence the level of genetic diversity and explain the strong genetic differentiation between the two *ex situ* collections. In summary, GBS is a useful method for characterizing genetic diversity in cauliflower genebank material and our results suggest that it may be useful to incorporate routine genotyping into accession management and seed regeneration to monitor the diversity present in *ex situ* collections and to reduce the loss of genetic diversity during seed regeneration.

## Introduction

The extent and type of genetic variation present in the germplasm of a crop is an important component of efficient breeding programs, because it provides useful information for the broadening of breeding pools, the utilization of heterosis and the selection of parental lines. Also, this information helps breeders to narrow the search for new alleles at loci of interest and assists in the identification of markers linked to desirable traits for introgression into new varieties [1]. An assessment of genetic diversity is also essential for the organization, conservation and use of genetic resources to develop strategies for optimal germplasm collection, evaluation and seed regeneration [2].

*Ex situ* conserved plant genetic resources (PGR) are plant genotypes that are stored in central storage facilities. PGR are utilized to improve modern cultivars by the introgression of new and exotic genetic variation into breeding pools (e.g., [3]). However, PGR often experience a loss of genetic diversity, stronger inbreeding depression (especially in outcrossing crops) and accumulation of deleterious alleles because of small population sizes of individual genebank accessions. These processes may negatively affect the success of *ex situ* conservation after several regeneration cycles [4, 5]. In addition, strong selection caused by adaptation to the seed regeneration environment may further reduce genetic variation.

Cauliflower (*Brassica oleracea* var. *botrytis*) is an important vegetable crop worldwide and a valuable component of a healthy diet because of a high content of glucosinolates with anticancer properties [6, 7]. Cauliflower and broccoli are currently cultivated worldwide on about 1.2 Mio hectares, with an annual production of over 21 Mio. tons [8]. Genetic diversity of cauliflower was analyzed with a diversity of marker systems like amplified polymorphic DNA (RAPD; [9]) or simple sequence repeats (SSRs; [10, 11]). These initial genotyping studies indicated that genetic diversity for cauliflower was limited [11–13]. More recently, whole genome resequencing revealed the genetic structure of cauliflower germplasm and identified genomic regions with low and high genetic diversity, respectively [14]. Whole genome sequencing approaches are still too expensive for large numbers of genebank accessions, or not necessary because smaller polymorphism numbers are sufficient for the analysis of diversity and genetic relationships. Reduced representation sequencing methods such as genotyping-by-sequencing (GBS) are a cost-effective alternative because these methods allow a high degree of multiplexing and tens of thousands of polymorphisms are identifed in a single reaction without the need of a reference sequence [15–17]. In the context of PGR, GBS was used to characterize the genetic variation of maize, sorghum and switchgrass with respect to their known ancestral history and geographical origin [18–20]. In Brassicaceae, GBS was used to analyse genetic diversity in yellow mustard [21].

The density of polymorphisms identified by reduced representation sequencing methods like GBS are sufficient to conduct genome-wide analyses of diversity and to construct core collections for further phenotyping or breeding. Our objective was to use GBS for assessing the genetic diversity and genetic relationship of randomly selected accessions of the USDA and IPK *ex situ* genebanks, which harbor large collections of cauliflower accessions. We also investigated whether imputation of missing genotypes improves diversity estimates and whether genetic and phenotypic diversity among cauliflower accessions are correlated. Cauliflower is a predominately outcrossing species whose pollination depends on insects. A self-incompatibility (SI) system prevents self-fertilization, but high variation in the extent of SI was reported in different landraces and varieties [22]. This variation has been used to select highly inbred and highly homozygous lines for breeding. Both SI and cytoplasmatic sterility (CMS) mechanisms are employed in hybrid breeding of modern varieties, which is now the predominant breeding method. For these reasons, genebank accessions of cauliflower may be highly variable in their genetic diversity. In the present study, we focused on characterising species-wide diversity and included only a single individual of each accession to maximise diversity of number of accessions given the resources available. We observed a high level of population structure among accessions, and in particular a strong genetic differentiation between accessions from the two genebanks.

## Materials and Methods

### Plant material

A total of 191 cauliflower accessions were randomly selected and ordered from the genebanks of the United States Department of Agriculture (USDA), USA and IPK Gatersleben, Germany. They represent 47% (100 of 212) of the USDA and 40% (91 of 227) of the IPK cauliflower accessions, respectively. We selected accessions from these two large genebanks because they harbor large collections of cauliflower that are expected to reflect the worldwide diversity of this vegetable crop. According to the passport information, the sample consists of traditional cultivars, breeding material, hybrids, unverified genotypes, collector material, commercial vegetable seeds and landraces (Table 1). Accessions originate from 26 countries and 11 accessions are of unknown geographic origin. In addition, a single plant of the wild type of *Brassica oleracea* was obtained from Heidelberg Botanic Garden and Herbarium (HEID), Germany. All accessions of this study including accession type and country of origin are listed in Table S1.

**Table 1.**
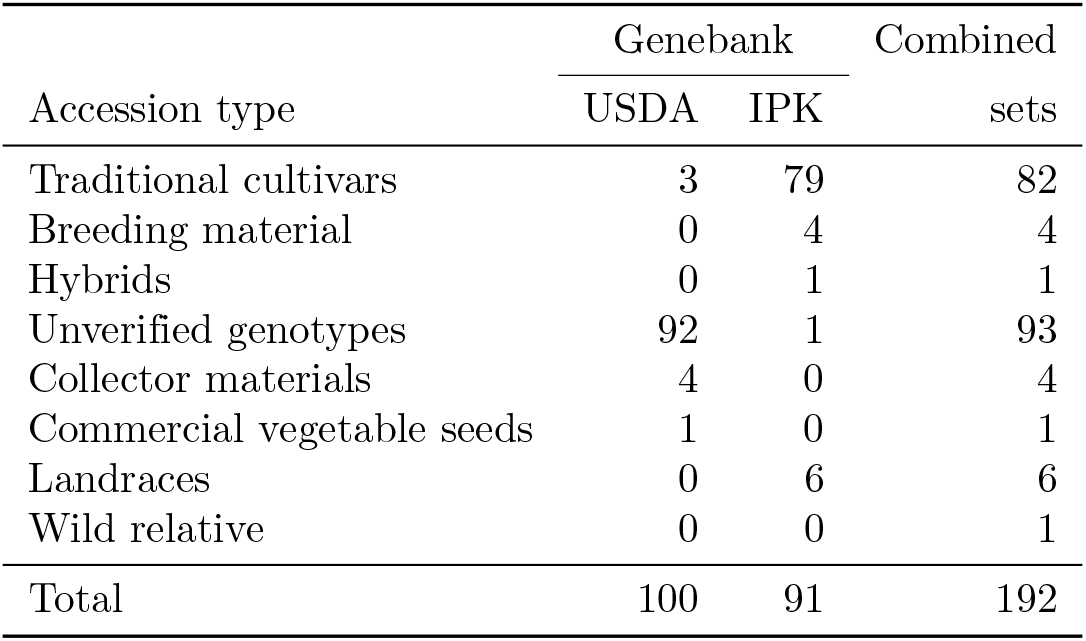
Counts of different types of genebank accessions genotyped.

### Field experiment and phenotypic measurements

All accessions were evaluated for six morphological traits at six environments consisting of two cultivation methods (organic and conventional) and three growing seasons (June 2011 and April/August 2012) in a randomized complete block design (RCBD) with two replicates in each environment [23]. Five random plants per plot were evaluated every three days for the following traits: curd width (cm), cluster width (cm), number of branches, length of the apical meristem (cm), length of the nearest branch to apical meristem (cm) and number of days from planting to appearance of the floral buds. Morphological traits were measured according to Lan and Patterson (2000) [24]. Phenotypic values were calculated as mean over locations and seasons.

### DNA extraction and genotyping by sequencing

Genomic DNA was extracted from leaf tissue three weeks after sowing in the green house from a single individual of each accession according to a standard CTAB protocol [25]. The quality and quantity of extracted DNA were checked with Nanodrop 2000c (Thermo Scientific), Qubit 2.0 Fluorometer (Life Technologies) and 3% agarose gel. The final concentration of each DNA sample was adjusted to 100 ng/ml for DNA digestion. GBS was carried out according to the protocol of Elshire et al. [15] with minor modifications. The DNA was digested with the ApeK1 restriction enzyme. A total of 96 barcodes were used, of which 64 barcodes were obtained with the web tool at http://www.deenabio.com/services/gbs-adapters and 32 barcodes were taken from [15]. All barcodes have an even distribution in length (4-8 nucleotides) and nucleotide composition of nucleotides at each position (Table S2). The 192 genotypes were divided into two libraries, each consisting of 96 genotypes. Before sequencing the GBS libraries, the distribution of fragment sizes were determined with an Agilent 2100 Bioanalyzer (Agilent Technologies, Santa Clara, CA) to verify that adapter dimers are absent and DNA fragments range between 170-350 bp [15]. The two libraries were sequenced on two lanes of an Illumina HiSeq1000 at the Kompetenzzentrum Fluoreszente Bioanalytik (KFB), Regensburg, Germany to produce 100 bp long paired-end reads.

### Sequence data analysis

Sequence reads were filtered for sequencing artifacts and low quality reads with custom Python scripts, bwa [26] and FastQC [27]. Reads mapping to the PhiX genome, which is used for calibration in Illumina sequencing, were identified and removed with bwa. All reads with ambiguous ‘N’ nucleotides and reads with low quality values were discarded. Remaining sequence reads were demultiplexed into separate files according to their barcodes. After removal of the barcode sequence and end-trimming, reads had a length of 88 bp. The pre-processed reads were then aligned to the genome of *Brassica oleracea* sp. *capitata* cabbage line 02-12 [28] with bwa. SNP calling was performed with SAMtools [29], bcfutils, vcfutils and custom Python scripts. The VCF file was parsed to retain SNP positions with a coverage of at least 30, whereby at least ten reads had to confirm the variant nucleotide. Positions not fulfilling these criteria were marked and considered as missing data. A distance matrix was calculated using the SNP data as input (Supplementary Note 1).

### Analysis of genetic structure and genetic diversity

The genetic structure of the sample was investigated by various methods for comparison. They included principal component analysis (PCA), as implemented in the R package adegenet [30]. Also, a PCA of six morphological traits was performed with the prcomp function in the stats R package [31] and a multivariate analysis of variation (MANOVA) with the manova function of R. For post-hoc tests of phenotypic differentiation, a single-factor ANOVA was calculated and a Bonferroni correction applied by applying a critical threshold of *p* = 0.05/6 = 0.0083 to test results. The correlation between genetic and phenotypic distances based on the PCA was determined with a Mantel test [32] in the ade4 R package [33]. Principal coordinate analysis (PCoA) based on pairwise *F_st_* values between genotypes calculated with R adegenet R package [30], was used for further analysis of population structure with the ape R package [34]. In addition, the genetic relationship among accessions was assessed with a neighbor-joining tree (NJ tree) based on a pairwise distance matrix with ape R package [34]. Population structure was inferred with ADMIXTURE [35]. The number of subpopulations analyzed ranged from *K* =1 − 10 and cross-validation was used to estimate the value of *K* which best fits the data [36]. An Analysis of Molecular Variance (AMOVA) was carried out with Arlequin v3.5.3.1 [37]. The extent of genetic differentiation (*F_st_*) between the two genebanks was estimated with the pairwise.fst function in the R package adegenet [30]. For each group of accessions from USDA and IPK genebanks, we calculated the observed and expected heterozygosities, *H_o_* and *H_exp_*, as well as the inbreeding coefficient (*F*) with adegenet. We also calculated percent polymorphic loci, %P, and nucleotide diversity, *π*, with the R package pegas [38]. To compare the effects of missing values and of genotype imputation on the population structure inference and genetic diversity estimates, we carried out the analyses with three data sets: 1) data without missing values, where all markers with missing values were excluded; 2) data with missing values, in which all markers with missing values were retained; and 3) imputed data, in which the missing values were imputed with fastPHASE [39].

### Detection of outlier SNPs

Outlier tests for highly differentiated SNPs between the sets from the two genebanks (USDA and IPK) were based on a coalescent-based simulation method [40] implemented in LOSITAN [41]. This method identifies putative targets genes that differentiated in response to selection based on the distributions of expected heterozygosity and *F_st_* values under an island model. LOSITAN was run in two steps: in the first step, an initial run with 50,000 simulations was performed with all SNPs and using the mean *F_st_*. After excluding a candidate subset of selected SNPs determined in the initial run, the distribution of neutral *F_st_* values were calculated. We also used Arlequin 3.5 [37] to detect outlier SNPs by accounting for a hierarchical genetic structure based on 10,000 simulations under a hierarchical island model. For both methods, SNPs outside the 99 and 1% confidence areas were identified as candidate genes potentially affected by directional and balancing selection, respectively. A further test of differentiation by selection was conducted with BayPass v2.1 [42] which is a fast implementation of the Bayenv2 algorithm [43].We first estimated the population covariance matrix Ω, which is a variance-covariance matrix of allele frequencies, for the two groups of accessions from each genebank was calculated from the final run of the MCMC after 100,000 iterations. We used 360 (25%) randomly selected SNP markers from the SNP dataset without missing values. To evaluate the robustness of this matrix, we repeated the calculation with five randomly drawn SNP sets and calculated the average correlation coefficient of estimated variance and covariance parameters from all 10 pairwise comparisons of matrices. The resulting average correlation coefficient was 0.99, which indicates a high convergence of Ω matrices based on different random SNP sets. The Ω matrix was then used to control for the genome-wide genetic relationship among groups in the calculation of the *X^T^X* statistics for each SNP by using 100,000 iterations of the MCMC. The *X^T^X* statistic is equivalent to the *F_st_* value as a measure of population differentiation.

### Data availability

Raw sequence data have been submitted to Short Read Archive (SRA) under accession number PRJEB8701. Aggregated data (e.g., SNP calls) and analysis scripts are available on the Zenodo data repository (DOI:10.5281/zenodo.887955).

## Results

### SNP identification by GBS analysis

To analyse genetic diversity in the sample we included one individual plant per genebank accession. The sequencing data of the 192 accessions consisted of 455 Mio. reads with a length of 100 bp. Quality filtering removed 5.2% of reads because they mapped to the PhiX genome or were of low quality, and read number per accession ranged from only 80 to 7.1 Mio with an average of 2.4 Mio reads. Eighteen accessions with less than 200,000 reads were excluded from further analysis because the proportion of missing data in these accessions was too high. A total of 133 Mio reads mapped to the *B. oleracea* reference genome (Figure 1). The percentage of mapped reads per genotype against *B. oleracea* ranged from 14% to 35% with an overall average of 29% (Table S3). Based on the mapping to the *Brassica oleracea* reference genome, 120,693 SNPs were detected in the remaining 174 samples (120,693 SNPs with missing data and 1,444 SNPs without missing data in any of the accessions). The mean percentage of missing data across all genotypes was 42% and values ranged from 19% to 77% per genotype. The number of SNPs and percentage of missing data per genotype are shown in Table S4.

**Fig 1.**
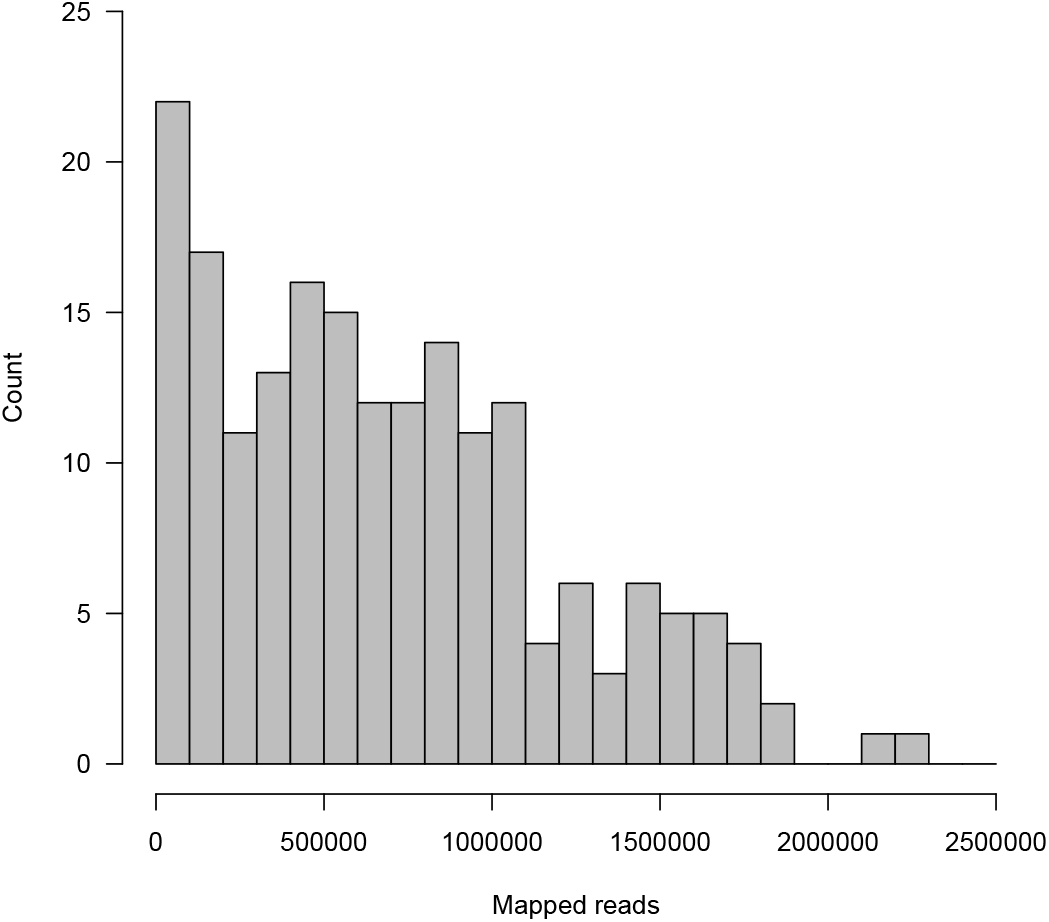
Histogram of read counts mapped to the reference genome.

### Analysis of genetic population structure

The genetic structure of the whole collection (*n* = 174) was analyzed with PCA, PCoA and ADMIXTURE. Here, we present the results for the set of 1,444 SNPs without missing data, but the same results were obtained with the other two data sets with missing or imputed data (in both cases, *n* = 120, 693), which are provided as Supplementary Information.

The marker-based PCA showed a clear differentiation between the two genebanks for the SNP data (Fig. 2A). The first two axes explained 21% of the overall variance and separated accessions from the USDA and IPK genebanks. The SNP data sets with missing and imputed values showed the same genetic structuring (Fig. S1).

**Fig 2.**
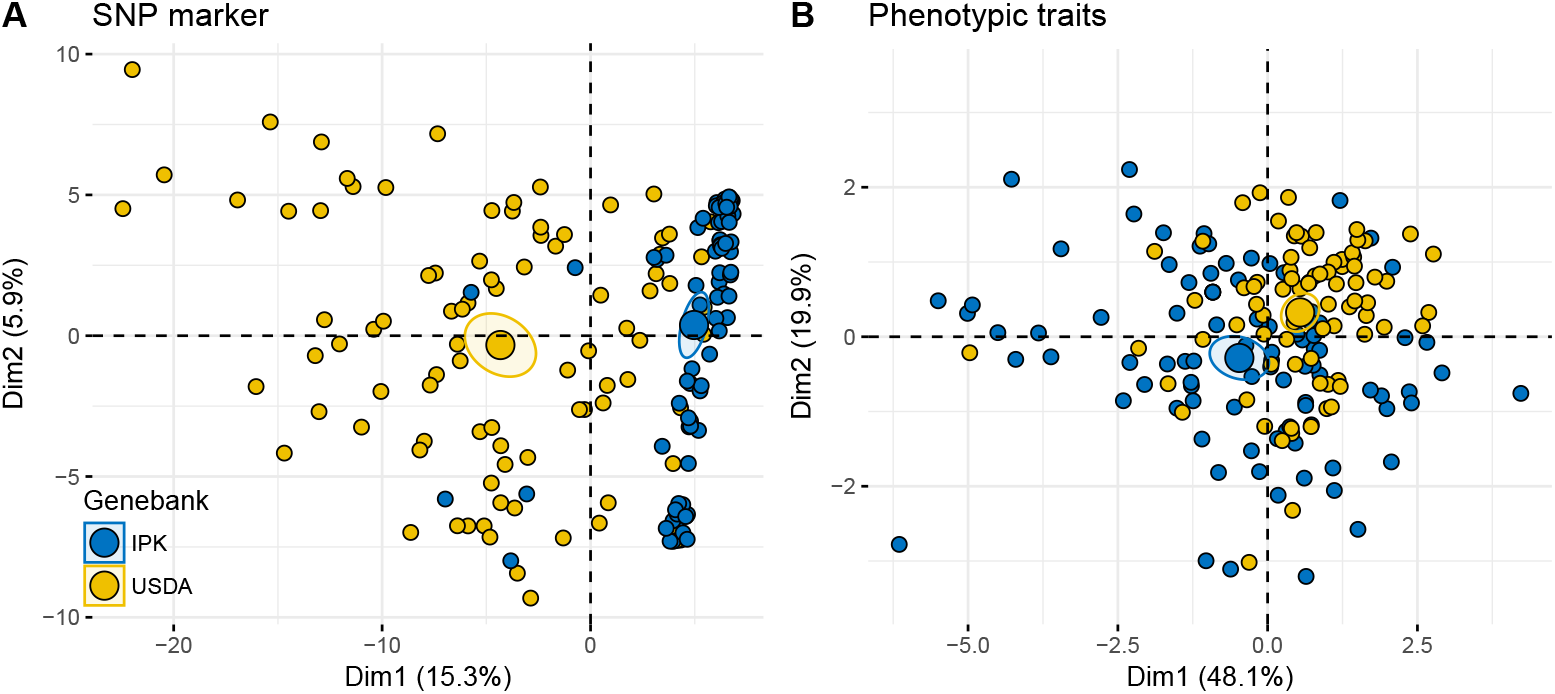
Principal component analysis (PCA) of SNP markers and phenotypic traits. (A) PCA of 1,444 GBS-derived SNPs without missing data in any of the accessions. (B) PCA of six phenotypic traits measured in two locations over three growing periods. In both PCA analyses, the two larger dots indicate the centroids and their confidence regions for the genebanks.

Further analyses confirmed the genetic differentiation of accessions and reveal that the largest proportion of genetic variance is explained by the difference between the two genebanks. A PCoA based on pairwise *F_st_* values separated the USDA from the IPK accessions on the first principal component axis, which explained 24% of the overall variance, whereas the second axis explained only 8% of the variance (Fig. S2). A neighbor joining (NJ) tree based on a pairwise distance matrix separated the 174 accessions into two distinct groups (Fig. 3A) representing the two genebanks. In both groups, accessions are not differentiated into well-supported subgroups that reflects the country of origin (Fig. 3B). The NJ trees based on the SNP data with missing or imputed data (120,693 SNPs in both data sets) confirm these results (Fig. S3).

**Fig 3.**
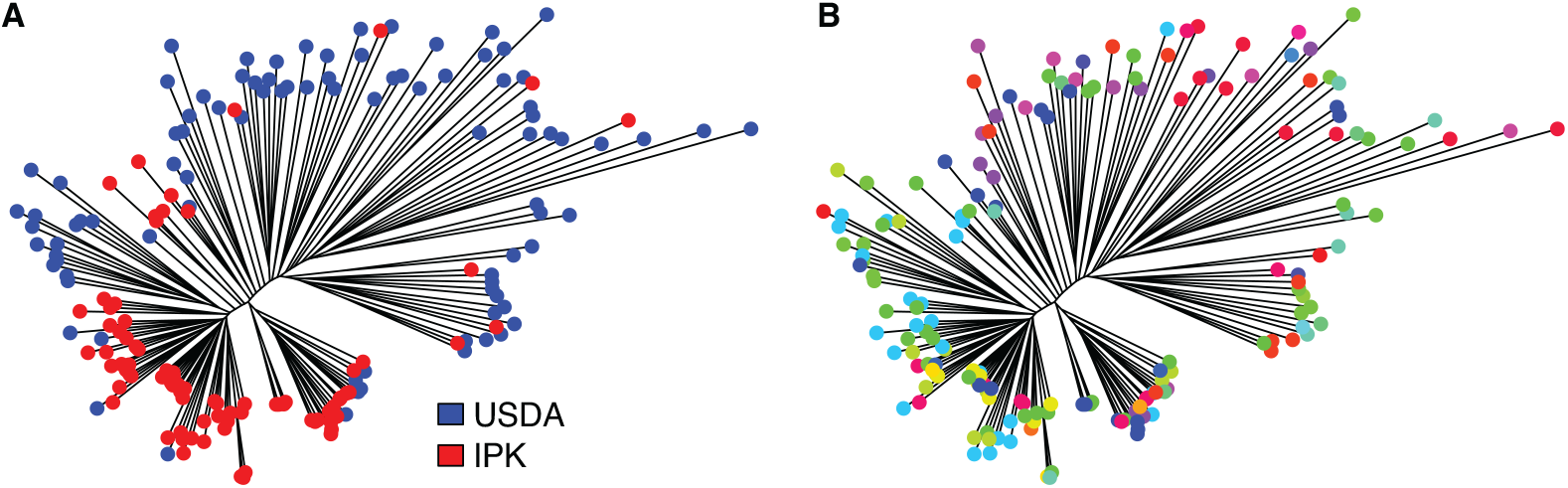
Neighbor joining tree of 174 cauliflower accessions. The NJ tree is based on the pairwise distance matrix using data without missing values (1,444 SNPs). (A) The two genebanks are represented as different colors. (B) Each country of origin of the original seeds as stated in the passport information is represented with a different color.

In addition to the differentiation between genebanks the previous analyses also indicate the presence of additional clusters. We therefore used ADMIXTURE to infer population structure and to estimate the number of genetic clusters that is most consistent with the data. For *K* = 2, two groups mainly differentiated between the two genebanks (Fig. 4, S4 and S5). Based on cross-validation, ADMIXTURE identified five genetically different clusters as most consistent with the GBS data without missing values With *K* = 5, the IPK accessions cluster into two distinct groups with some degree of admixture, whereas the USDA accessions do not form distinct clusters and show a high level of admixture (Figure 4).

**Fig 4.**
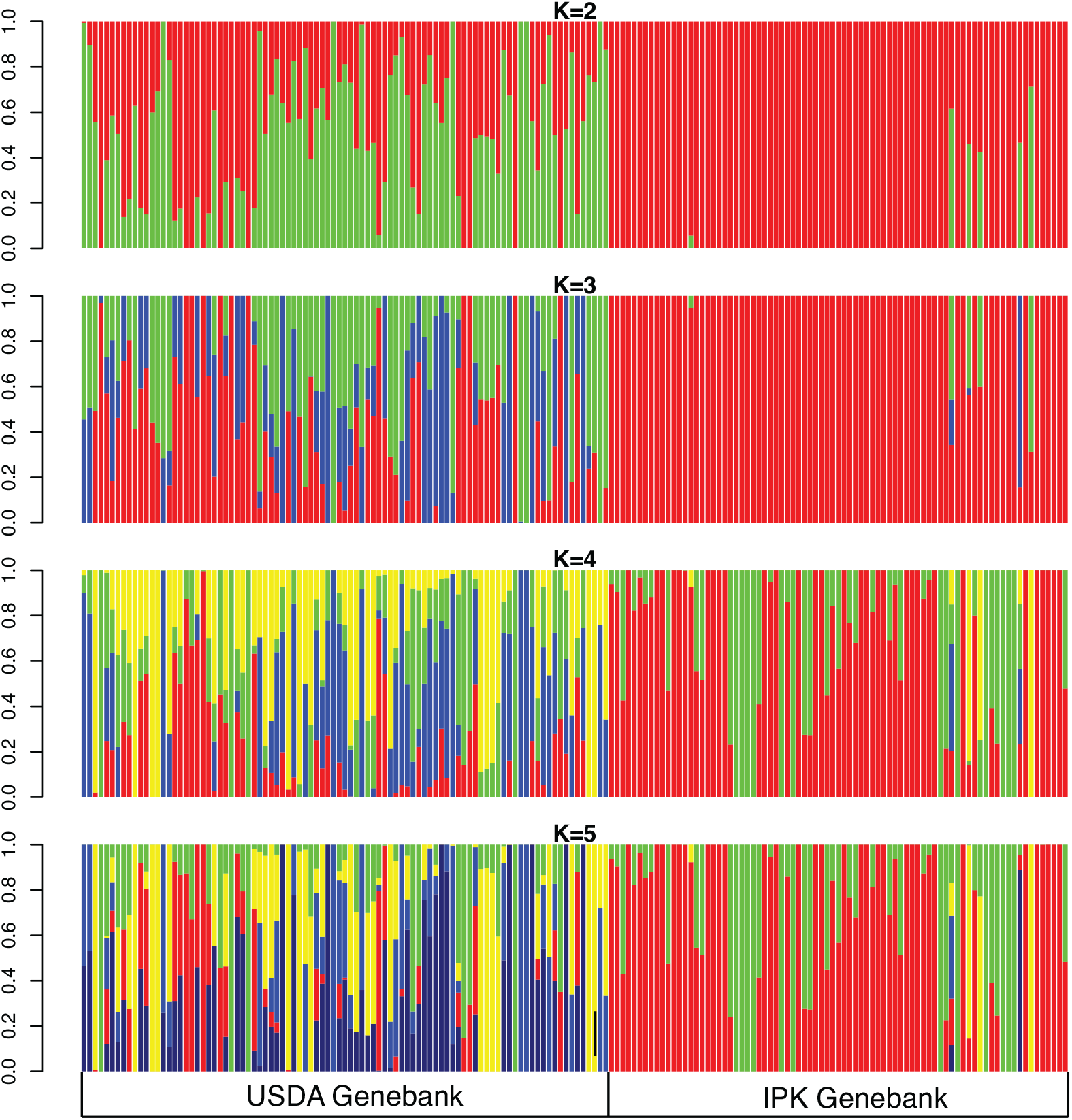
Genetic structure of 174 cauliflower genotypes inferred with ADMIXTURE. The number of predefined clusters ranged from *K* = 2 − 6 and was inferred using SNPs without missing values (*n* = 1, 444).

Despite the clear genetic differentiation between the two genebanks, a very large proportion of genetic variation (90%) segregates within rather than between genebanks (5%; AMOVA, *p* < 0.001; Tables 2, S5 and S6). This is consistent with a low overall genetic differentiation between the two genebanks (*F_st_* = 0.029). The mean pairwise *F_st_* of accessions within each genebank was 0.301 for the USDA and 0.160 for the IPK accessions (estimated from 1,444 SNPs without missing values), showing that the USDA accessions are more differentiated from each other than the IPK accessions.

**Table 2.** Analysis of molecular variance (AMOVA) of different groups based on SNP data without missing values.

| Level of variation | Sum of Squares | Variance component | % variance explained | P-value |
| --- | --- | --- | --- | --- |
| Among groups (genebanks) | 1,032 | 5.43 | 6 | <0.001 |
| Within groups | 31,448 | 91 | 94 | <0.001 |
| Total | 32,489 | 96 |  |  |

### Levels of genetic diversity

For a further comparison between accessions from the two genebanks, we calculated various genetic diversity parameters (Table 3). Values for expected heterozygosity, observed heterozygosity, percentage of polymorphic SNPs and nucleotide diversity were all larger in the USDA than in the IPK accessions. Accessions showed a high inbreeding coefficient (*F* > 0.5), which did not differ between both genebanks.

**Table 3.** Measures of diversity within two genebanks based three different data sets. Sample size of USDA accessions, *n* = 93 and of IPK accession, *n* = 81.

| Statistic | SNPs without missing values<br>(1,444 SNPs) |  | SNPs with missing values<br>(120,693 SNPs) |  | Imputed SNPs<br>(120,693 SNPs) |  |
| --- | --- | --- | --- | --- | --- | --- |
|  | USDA | IPK | USDA | IPK | USDA | IPK |
| $H_e$ | 0.157 | 0.088 | 0.214 | 0.149 | 0.201 | 0.107 |
| $H_o$ | 0.167 | 0.127 | 0.419 | 0.339 | 0.094 | 0.077 |
| $F$ | 0.501 | 0.500 | 0.501 | 0.501 | 0.495 | 0.507 |
| $\pi$ | 0.158 | 0.089 | 0.157 | 0.095 | 0.202 | 0.110 |

$H_e$ = Expected heterozygosity, $H_o$ = Observed heterozygosity, $F$ = Inbreeding coefficient, $\pi$ = nucleotide diversity.

### Identification of highly differentiated outlier SNPs

The strong genetic clustering of accessions from the two genebanks despite a low overall *F_ST_* value suggests that a small number of SNPs are responsible for the differentiation. To identify polymorphisms whose allele frequencies differ between the two genebank collections, we performed outlier tests using the GBS data without missing values (1,444 SNPs). LOSITAN identified 182 (12.6%) SNPs as outliers (Table S7) based on *F_st_* values between the two populations (FDR < 0.1). Arlequin identified only 79 (5.5%) SNPs as outliers (Table S8). Since no *p*-values can be calculated for the *X^T^X* statistic, we ordered SNPs by the rank order of *X^T^X* values, as suggested by [43] to detect strongly differentiated SNPs. A total of 73 SNPs were among the top 5% (Table S9). The Venn diagram in Figure 5 shows little overlap of highly differentiated SNPs detected between pairs of methods, and only one SNP was identifed as an outlier by all three methods.

**Fig 5.**
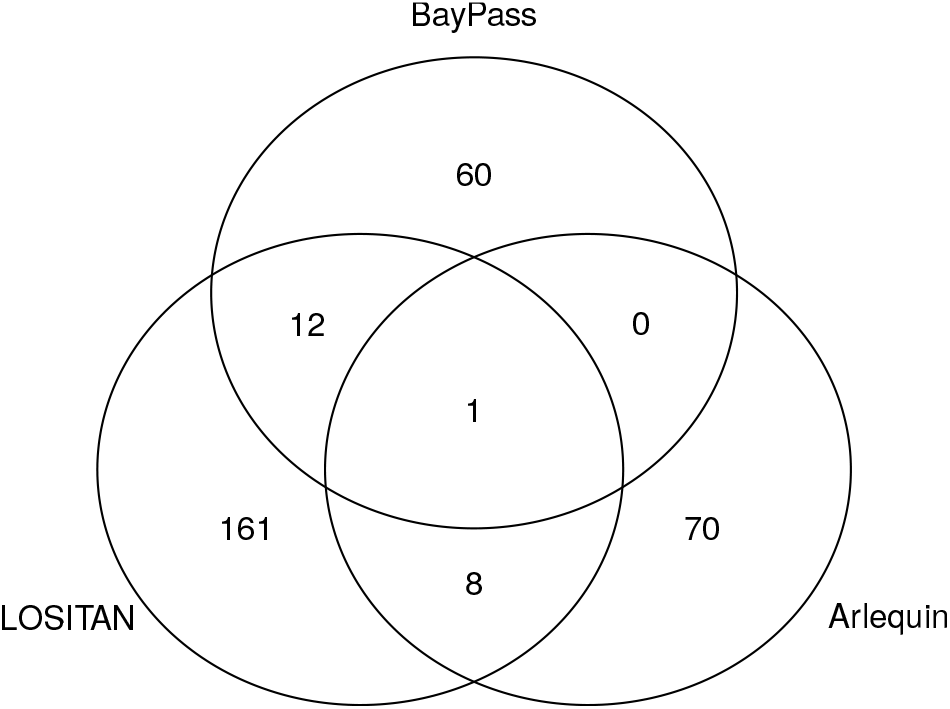
Overlap between outlier SNPs. Venn diagram of all significantly differentiated SNPs detected with Arlequin (*p* < 0.05), LOSITAN (*p* < 0.05) and BayPass (top 5% X^*T*^X values).

### Phenotypic differentiation between genebanks

Since the genetic analysis indicated a strong differentiation of accessions from the two genebanks, we also investigated their phenotypic differentiation. A PCA of phenotypic traits indicated a weaker differentiation between USDA and IPK accessions (Fig. 2B) than the genotyping data, although the centroids of each cluster are clearly differentiated. The IPK accessions clustered more strongly than the USDA accessions which indicates a lower phenotypic variation and is also consistent with the genotyping data. The lower phenotypic than genetic differentiation between the genebanks reflects genotype x environment (GxE) interactions of phenotypic traits as inferred in a previous study [23]. The first principal component explained almost half of phenotypic variation in the total sample and the second about 20% (Figure 6A). The variable correlation plot indicates that the curd related traits are strongly related to the first principal component, whereas days to bolting (flowering time) shows a very high correlation with the second principal component (Figure 6B).

**Fig 6.**
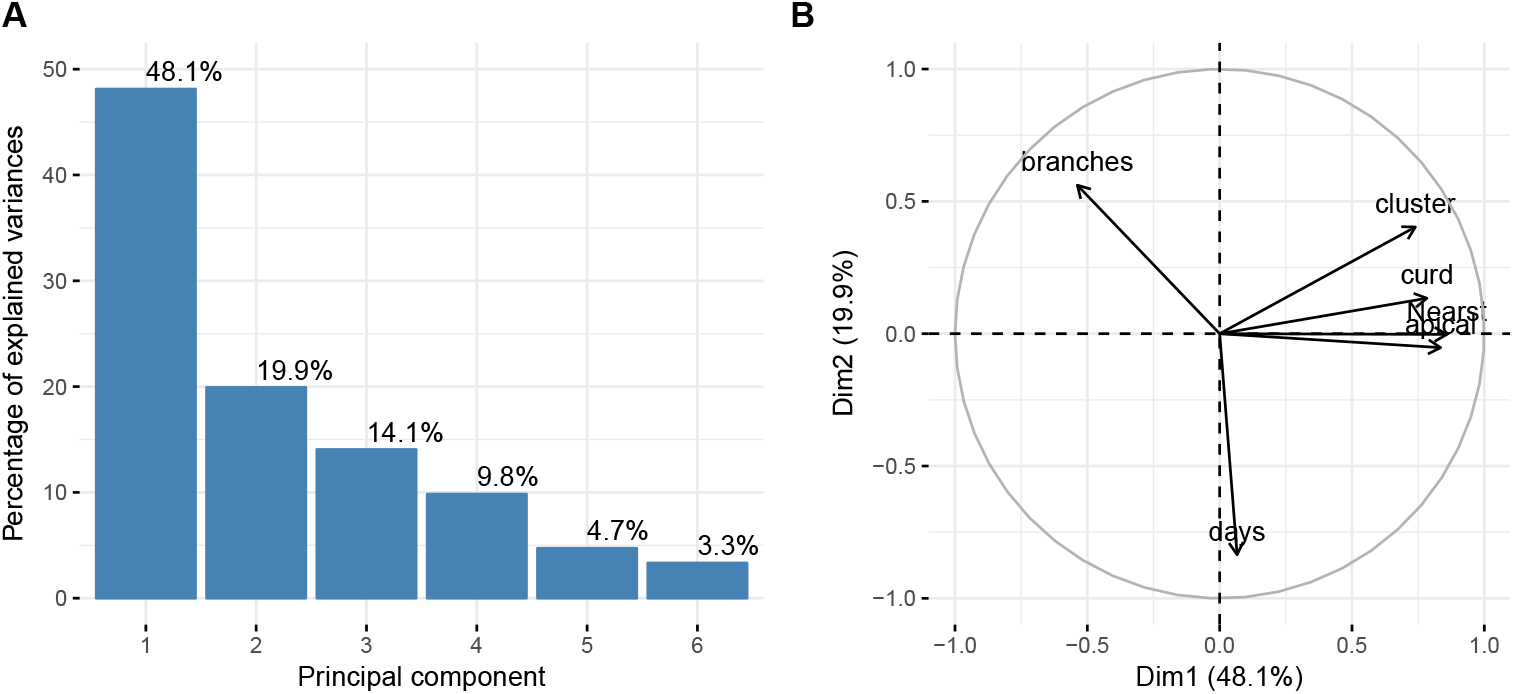
Parameters of PCA with phenotypic variation. (A) Percent variance explained by each of the six principal components. (B) Variable correlation plot on the first two principal components. The quality of traits on the PC map is expressed as distance between traits and the origin. Traits that are distant from the origin are well represented on the factor map.

To test which traits are significantly differentiated between genebanks, we used MANOVA for comparison because some traits are strongly correlated (Figure S6). The strongest correlation is between length of nearest branch to apical meristem and length of apical meristen (*r* = 0.798, *p* = 10^−12^). The phenotypic variance was different between accessions from both genebank, and the MANOVA with genebank as grouping factor strongly supported the phenotypic differentiation (Pillai test statistic: 0.3158, *p* < 10^−11^). A post-hoc analysis of single-factor ANOVA with Bonferroni correction revealed that only the traits curd width (F = 15.4, *p* = 0.0001254) and cluster width (*F* = 67.8, *p* < 10^−14^) differed between genebanks.

Finally, a Mantel test showed a positive correlation of phenotypic and genetic similiarities between pairs of accessions (*r* = 0.291, *p* < 0.001).

## Discussion

### Assessment of genetic diversity by GBS

Previously, the genetic diversity of cauliflower was assessed with different types of markers and based on small data sets. This limitation can be overcome by sequencing-based methods like GBS [15]. Although we generated a large number of raw reads, a substantial proportion of reads (71% on average) did not align to the *B. oleracea* reference genome, similar to what has been observed in maize [18]. Such a high proportion of unmapped reads may result from the evolutionary divergence between the subspecies *B. oleracea* spp. *capitata* (cabbage) from which the reference genome was produced and cauliflower, *B. oleracea* var. *botrytis*. This interpretation is confirmed by the pan-genome analysis of *B. oleracea* cultivars [14, 44]. Another explanation is that the reference genome is still incomplete. The proportion of matching reads may be also influenced by a limited sensitivity of the Burrows-Wheeler Alignment (BWA) algorithm or a high proportion of presence/absence variation (PAV) [18].

GBS was used for a wide range of species and is an effective method for generating tens of thousands SNP markers [15, 17], but the high proportion of missing data, which in our study varied between 19% and 77% per accessions, is a major disadvantage [15]. Possible solutions to reduce the proportion of missing data include the sequencing to higher read depths [16] or the imputation of missing values, which we pursued in this study. A comparison of diversity estimates obtained with the three SNP sets indicates that estimates of observed heterozygosity, *H_o_*, are influenced by missing data, whereas estimates of the inbreeding coefficient and nucleotide diversity, *π* are very similar (Table 3). Therefore, genome-wide parameters like diversity or genetic structure can be estimated with or without missing data. Due to the rapidly decreasing sequencing costs and protocols for low-cost preparation of whole genome sequencing libraries [45], low coverage genome sequencing is rapidly becoming the method of choice for characterizing the genomic diversity of species with moderate genome sizes such as *B. oleracea*. The correlation of phenotypic and genotypic distance as indicated by the Mantel test indicates that genotyping can be used to assemble phenotypically diverse core collections for further evaluation.

### Patterns and causes of genetic structure in cauliflower accessions

Despite the high proportion of missing data, GBS allowed to analyze genetic diversity and population structure in *B. oleracea* genebank accessions. Our sample of cauliflower accessions form groups that do not reflect their geographic origin but the seed source (i.e., *ex situ* genebank). A geographic population structure was found in previous survey of cauliflower cultivars [9] despite a smaller sample size than our study. Furthermore, patterns of genetic diversity of switchgrass, maize and sorghum genebank accessions obtained with GBS was consistent with the ancestral history, morphological types and geographic distribution of these crops [18–20]. The clustering and the different levels of genetic diversity between genebanks suggest that other factors than geographic origin determine the genetic relationship of accessions.

First, the accession types (landraces vs. cultivars) and collection dates may result from different collection strategies between genebanks that may be responsible for the observed clustering of accessions. For example, the USDA sample includes a much higher proportion of unverified material than the IPK sample which mostly consist of modern varieties. Passport data suggest that many accessions in the ‘unverified’ group of USDA and the ‘cultivar’ group of IPK reflect common cultivars (or possibly landraces) that were commercially available during the time of collection. Five accessions in the IPK set differed markedly from the other IPK accessions in all analyses and they clustered together with the set of USDA accessions. They consist of four landraces and one hybrid (Table S1), supporting the notion that the USDA genbank included more sources of germplasm that resulted in a larger collection of genetically diverse landraces (which were labeled as ‘unverified’) than the IPK genbank. The significant difference in average collection date between both genebanks may further contribute to their differentiation. According to the passport data, the USDA accessions are on average more than 15 years older than the IPK accessions. The USDA accessions were collected between 1948 to 1981 with an average of 1959, whereas the IPK accessions were collected between 1957 to 2002 with an average of 1974. For this reason, the observed population differentiation may also be caused by various process such as breeding progress, seed management strategies, inbreeding or genetic drift of *B. oleracea* germplasm. Unfortunately, limited passport information did not allow us to account for the breeding history and relationships among varieties as co-variate in the population structure analysis and to disentangle the effects collection date and geographic origin on diversity estimates. In addition, the absence of a strong geographic structure may result from a combination of low genetic diversity in cauliflower [11, 12], an exchange of seeds over large distance in historical time, and a high level of gene flow due to outcrossing with other varieties [46]. On the other hand, the lower genetic diversity of the IPK relative to the USDA accessions may be caused by a higher proportion of modern cultivars, which have a narrower genetic basis resulting from modern breeding methods. These different processes are difficult if not impossible to reconstruct and confound the analysis of genetic diversity.

A second explanation for differentiation between genebanks are seed regeneration procedures which may affect genetic diversity [47–50]. In several species, *ex situ* conserved genetic resources had a lower diversity than *in situ* conserved populations [51–53] or historical material [46]. A reduction in the diversity of *ex situ* genebank material is mainly caused by a small number of individuals per accession that are usually conserved. Such accessions are exposed to genetic bottlenecks, inbreeding depression, the accumulation of mildly deleterious mutations and a loss of genetic diversity by random drift [6, 52, 54, 55]. A high overall inbreeding coefficient of >0.5 for all accessions estimated from the GBS data suggest an impact of small population size on genetic diversity. The inbreeding coefficient depends on the breeding history of the material before inclusion into the genbank and the seed propagation protocols. The seed regeneration procedures at the USDA and IPK genebanks likely did not contribute much to the observed differentiation. Seed regeneration at the USDA genebank is carried out in 12 x 24 ft cages (corresponding to 26.8 m^2^) with mesh covers to prevent cross-pollination by insects and a population size of at least 100 plants per accession. At IPK, cauliflower accessions are cultivated in small glass houses containing other species as well. The total area is about 6 m^2^ and population size is 20-25 plants. At IPK, seeds are regenerated after 20 years and at USDA after 15 years, which on average corresponds to 1.5 and 4 regeneration cycles for the material included in this study. Using the formula for calculating the expected decay in heterozygosity, *H_t_* = *H*_0_ × (1 − 1/*N_e_*)^*t*^ [56], the expected relative decay in heterozygosity of USDA genebank accessions is Δ*H* = (1 − 1/100)^4^ = 0.96 under the assumption that on average each accession has undergone 4 regeneration cycles, and of IPK accessions Δ*H* = (1 − 1/20)^1.5^ = 0.93. Although a more rapid decay in heterozygosity is expected in the IPK collection, the difference is rather small.

Natural and/or artificial selection during seed propagation of genebank accessions may also impact genetic diversity and differentiation in *ex situ* genebanks. The cauliflower accessions were collected in different regions of the world, and selection of genes controlling photoperiod sensitivity, flowering time, and other traits caused by local adaptation to the propagation environment may have occured. The effect of selection on fitness and genetic diversity can be quite strong. A significant correlation between genetic and phenotypic distance shown by the Mantel test and a high heritability of traits like flowering time [23] indicate a strong phenotype-genotype relationship that facilitates a rapid evolutionary response to selection. During seed regeneration, pollination is managed with commercial pollinators like bumblebees, but the reproductive success is not closely monitored and the effect of selection on the relationship structure is unknown. To test the potential impact of selection, we used three outlier tests to identify highly differentiated SNPs. Out of 1,444 tested SNPs without missing data, 12.5% (LOSITAN), 5.5% (Arlequin) and 5% (BayPass) were classified as highly differentiated between the two sets of accessions. The three methods differ in their approach to control for population structure and kinship to reduce the proportion of false positives. Shimada *et al*. [57] suggested to consider only SNPs that were identified by more than one method as true outliers, and such an approach was further confirmed in a simulation study of non-equilibrium populations [58]. Hence, in a comparison of outliers identified in our data (Figure 5), we identified only 0.8% (12 out of 1,444 SNPs) of SNPs as outliers by LOSITAN and BayPass, 0.6% (0) by LOSITAN and Arlequin, 0% (0) of SNPs by BayPass and Arlequin, and only a single SNP by all three methods. Overall, this is a small proportion (<1%) of the total number of SNPs tested. In conclusion, if natural or unintentional artificial selection during seed regeneration contribute to genetic differentiation, it may either weak selection, affect only few genomic regions or occur in regions that were not tagged by the SNPs of this study. The identification of strongly differentiated SNPs rests on the assumption that both collections derive from the same ancestral population, which likely is not true for our sample. The comparison of allele frequencies of accessions over seed regeneration cycles with higher marker densities is a more powerful approach to detect the effect of selection on *ex situ* genebank material, and facilitates the close monitoring of allele frequency changes and a better management of *ex situ* germplasm collections.

### Characterizing genebank accessions with GBS

A major advantage of GBS and related methods is their applicability to any species. These methods do not entail setup costs like SNP arrays and do not cost much per individual genotype, but provide sufficient power for genome-wide analyses of population structure and genetic relationships. On the other hand, GBS has a high proportion of missing data that may reduce the power for correct estimation of population parameters. Data imputation was suggested as a solution because it can be accurate and then increase the quality of genomic selection or association mapping [59, 60]. In our study, however, a comparison between three GBS-derived data sets consisting of SNPs without missing values, SNPs with missing values, and imputed SNPs revealed only a minor effect of missing data and data imputation on the ability to infer the population structure, although diversity estimates differed significantly between imputed and non-imputed data (Table 2). This result confirms a previous study [61] in which the estimation of heterozygosity and inbreeding coefficients was less accurate with a high proportion of missing data and estimation biases were much smaller for data sets with missing values than for imputed data sets. Furthermore, the density of GBS-derived markers are frequently too low to detect footprints of selection [62] caused by the different history of genebank collections or ongoing selection during seed regeneration. The correlation of genetic and phenotypic differentiation of collections (Figure 7) indicates that GBS is a highly suitable approach for defining core collections and sets of genetically differentiated genebank accessions that are further used for whole genome sequencing, phenotypic characterization or the establishment of (pre-)breeding populations.

**Fig 7.**
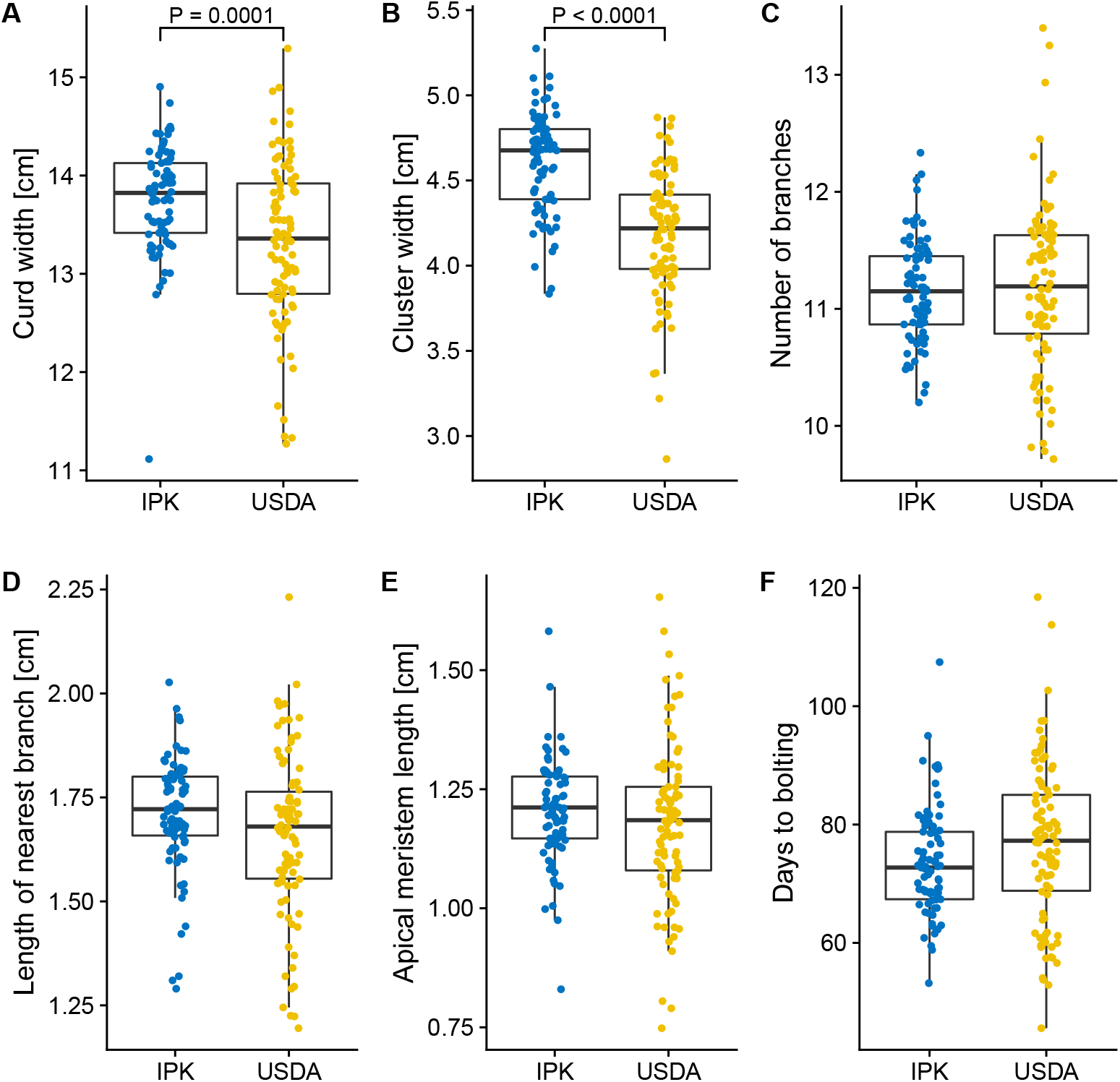
Box plots of six curd-related traits in accessions grouped by seed source (USDA and IPK genebanks). Significant differences between the two genebanks as observed from a single-factor ANOVA with Bonferroni correction of *p*-values. The *p*-values for the two traits with significant differences are shown. Individual phenotype values are averages of 6 environments (2 locations and 3 growing seasons each). The upper and lower hinges correspond to the first and third quartiles (the 25th and 75th percentiles), and the whiskers extend to the largest value no further than 1.5 * IQR (inter-quartile range), or distance between the first and third quartiles. Phenotypic data are from [23].

## Conclusions

Our study outlined the usefulness of GBS to characterize the genetic diversity of genebank accessions of a minor crop like cauliflower. A key result was the strong differentiation of genetic diversity between the two genebanks which most likely reflects the different collection histories of the two genebanks. Due to a lack of detail in the passport information, factors influencing genetic diversity like sampling strategy, regeneration procedures and selection during regeneration could not be well reconstructed, although the type of accessions included (landraces vs. cultivars) likely has a strong influence. The low cost of GBS and low-coverage genome sequencing suggest that a lack of passport information can be substituted by high-resolution genotyping and suitable analysis methods to characterize the diversity of germplasm from different sources. This facilitates the exchange of material between genebanks and the construction of core collections that harbor a high proportion of species-wide genetic and phenotypic diversity for a more efficient utilization of plant genetic resources [63]. GBS-derived polymorphisms may facilitate an exchange of germplasm between genebanks, but this requires an infrastructure for genomic data management similar to the information system already in place for passport data. Our work also demonstrated monitoring of genetic diversity during seed regeneration allows to manage diversity within accessions to mitigate some disadvantages of small population sizes of *ex situ* conserved plant genetic resources, in particular for outbreeding crops such as cauliflower.

## Supporting information

**S1 Fig. Comparison of the principal component analyses (PCA) of 174 cauliflower accessions.** (A) PCA based on the dataset with missing data and (B) using genotypes imputed with fastPHASE.

**S2 Fig. PCoA of pairwise** *F_st_* **values between individuals**.

**S3 Fig. Neighbor-joining tree of 174 accessions.** The tree is based on the pairwise distance matrix. (A) and (B) Analysis data with missing values. (C) and (D) analysis with imputed data. (A) and (C) Accessions are represented by different colors according to the genebank or origin. (B) and (D) Accessions are represented according to the country of origin.

**S4 Fig. Population structure analysis of 174 cauliflower accessions with ADMIXTURE based on SNPs with missing data**.

**S5 Fig. Population structure analysis of 174 cauliflower accessions with ADMIXTURE based on SNPs with imputed data**.

**S6 Fig. Correlations between six phenotypic traits of cauliflower**.

**S1 File. Calculation of the distance matrix**.

**S1 Table. Numbers, accession ID, accessions names, gene bank source and country of origin of accessions included in this study**.

**S2 Table. GBS Barcode IDs**.

**S3 Table. Number of raw read, mapped reads and percentage of mapped reads per accession**.

**S4 Table. Number of SNPs and percent of missing data per accession**.

**S5 Table. Analysis of molecular variance (AMOVA) of different groups based on SNP data with missing values**.

**S6 Table. Analysis of molecular variance (AMOVA) of different groups based on SNP data with imputed values**.

**S7 Table. Outlier SNPs** (*p* < 0.05) **among 1,444 SNPs without missing data identified with LOSITAN**.

**S8 Table. Outlier SNPs** (*p* < 0.05) **among 1,444 SNPs without missing data identified with Arlequin**.

**S9 Table. Top 5% of SNPs based on** *X^T^X* **values calculated with BayPass**.

## Acknowledgements

Keygene N. V. owns patents and patent applications protecting its Sequence Based Genotyping Technologies. We express our thanks to Fabian Freund, Christian Lampei, Dounia Saleh, Torsten Günther, Patrick Thorwarth and Linda Homann for discussions on data analysis methods. Also, we thank the USDA and IPK genebanks for prividing seed stocks. We thank Marcus Koch, Heidelberg Botanic Garden and Herbarium HEID, for seeds of the wild ancestor of *Brassica oleraceae*. This work was supported by a GERLS Fellowship of the DAAD to E. Y. and the F. W. Schnell Endowed Professorship of the Stifterverband für Deutsche Wissenschaft to K. J. S.

